# Twinfilin is a non-processive depolymerase which synergizes with formin to dramatically accelerate actin filament uncapping by 300-fold

**DOI:** 10.1101/2024.07.19.604324

**Authors:** Vishal Reddy, Ankita Arya, Shashank Shekhar

## Abstract

Cellular actin networks assemble by actin filament elongation at barbed ends and are thought to disassemble primarily by depolymerization at filament pointed ends. Contrary to this conventional understanding of actin dynamics, twinfilin was recently shown to promote barbed-end depolymerization. Twinfilin has additionally been suggested to sequester monomers and cap as well as uncap filament barbed ends. As a result, the exact mechanisms by which twinfilin affects barbed-end dynamics remain controversial. Using multicolor single-molecule microscopy, we show that both mouse and yeast twinfilin are non-processive depolymerases that interact only transiently with barbed ends (∼0.2-0.5 s). Each twinfilin binding event, on average, results in the removal of one or two actin subunits. At CP-capped barbed ends, twinfilin synergizes with formin to accelerate uncapping by up to ∼320-fold. We find that uncapping by twinfilin, alone and together with formin, depends on the nucleotide state of the filament, with the two proteins causing a much more modest enhancement of uncapping of newly assembled filaments. Our study thus establishes twinfilin as a multifunctional barbed-end binding protein capable of non-processively depolymerizing, transiently capping, and synergizing with formin to rapidly uncap actin filament barbed ends.

## Introduction

The dynamic actin cytoskeleton is a key determinant of cell shape and polarity, and plays an essential role in processes such as cell division and motility [1]. Traditionally, cellular actin dynamics have been thought to be governed by treadmilling, where actin filaments primarily elongate at their barbed ends [2, 3]. The filament barbed ends then get capped and the actin subunits in the filament age through ATP hydrolysis. Subsequent release of the gamma phosphate converts the ADP-P_i_ actin subunits to ADP-actin subunits [4, 5], which then get released back into the actin monomer pool via depolymerization from pointed ends of actin filaments [6].

Twinfilin, a member of the ADF/Cofilin-H protein family, has been a notable exception to the treadmilling dogma. Twinfilin was initially discovered as a G-actin binding protein that sequestered actin monomers in a 1:1 complex and prevented their assembly into filaments [7, 8]. Twinfilin localized to cortical actin patches in budding yeast and its deletion was synthetically lethal with a cofilin mutant (cof1-22) which displayed reduced actin binding and disassembly [7, 9]. Together, these observations suggested that like cofilin, twinfilin might also play a role in actin disassembly. In addition to binding G-actin, twinfilin also binds actin filaments at their barbed ends [10]. Nevertheless, how barbed-end interactions of twinfilin influence filament dynamics and the exact mechanism of this interaction have remained controversial. While two early studies suggested twinfilin capped barbed ends [10, 11], a later study instead suggested that twinfilin was a processive barbed-end depolymerase [12]. Additionally, we recently showed that twinfilin-mediated depolymerization is highly sensitive to filament age [13]. While it accelerates depolymerization of newly-assembled ADP-P_i_ filaments it reduces the rate of depolymerization of aged ADP filaments. Notably, contrary to the previous suggestion of twinfilin being a processive depolymerase, a recent structural study instead suggested that the it may act as a transient capper and a non-processive depolymerase, dissociating from the barbed-end along with terminal actin subunits [14].

We employed single-molecule microscopy to address these open questions. We find that fluorescently labelled twinfilin transiently associates with barbed ends of both newly-assembled and aged actin filaments. Twinfilin molecules remain bound to ADP barbed ends for about twice as long as ADP-P_i_ ends. Our analysis indicates that individual twinfilin binding events cause removal of one or two actin subunits at a time, thus suggesting that twinfilin might not be a processive depolymerase.

In addition to free barbed ends, we also investigated twinfilin’s effects on capping protein (CP)-bound filament ends. CP displays a very high affinity for barbed ends (*K_d_* = 0.1 nM) and in absence of other factors, spontaneous uncapping is a very slow process [15]. On average, a CP-capped barbed end takes about 30 minutes to uncap (*k_off_* = 3 – 4 x 10^-4^ s^-1^) [15–17]. Notwithstanding CP’s slow dissociation kinetics *in vitro*, CP displays rapid turnover *in vivo* (*k_off_* = 0.14 – 0.58 s^-1^) [18, 19]. This discrepancy between *in vivo* and *in vitro* uncapping rates suggests that additional proteins might participate in uncapping in cells. Twinfilin directly binds CP via its CPI-motif in its C-terminal “tail” domain and its localization to yeast cortical actin patches gets disrupted upon deletion of genes encoding for capping protein [20, 21]. Through this interaction, twinfilin accelerates dissociation of CP from barbed ends [22]. Twinfilin can thus potentially generate free barbed ends for its depolymerizing activity. Nevertheless, even at saturating concentrations, twinfilin on its own is only a modest uncapper and causes a sixfold acceleration in uncapping [22]. As a result, twinfilin-mediated uncapping remains orders of magnitude slower than CP dissociation rates *in vivo* (*k_off_* = 0.14 – 0.58 s^-1^) [18, 19], suggesting that twinfilin might synergize with other factors to accelerate uncapping.

Formin promotes elongation of actin filaments from profilin-bound actin monomers [23, 24]. Formin has also been shown to have an uncapping effect via the formation of a formin-CP ‘decision complex’ at barbed ends [25, 26]. Through this mechanism, formin enhances the dissociation rate of CP by ∼tenfold [26]. More recently, it was shown that twinfilin can also join formin and CP at the barbed end to form a three-protein elongator-capper-depolymerase complex in which each protein directly interacts with terminal actin subunits [27]. In light of these discoveries, we sought to determine if formin and twinfilin could collaborate in promoting uncapping. Our data show that these two proteins synergize to uncap barbed ends of ADP actin filaments and promote a ∼318-fold increase over spontaneous uncapping which is approximate twice as fast as that of CARMIL (∼180-fold) [17]. In our experiments, we observed a maximal uncapping rate of ∼0.1 s^-1^ (at 200 nM mDia1 and 5 µM mTwf1) which is close to the CP dissociation rate inferred from *in vivo* experiments (*k_off_* = 0.14 – 0.58 s^-1^) [18, 19]. Importantly, we found that uncapping was highly sensitive to the nucleotide state of actin subunits in the filament. In control reactions as well as in presence of formin and twinfilin (alone and together), ADP filaments uncapped significantly faster than ADP-P_i_ filaments. Interestingly, while the two proteins synergize to uncap ADP filaments, their effect is only additive on ADP-P_i_ filaments. Our results thus show that newly-assembled ADP-P_i_ filaments are more resistant to uncapping, both spontaneous as well as in presence of twinfilin and formin.

## Results

### Twinfilin interacts transiently with filament barbed ends

We first investigated whether mammalian twinfilin can function as a processive depolymerase of actin filament barbed ends. Mouse twinfilin-1 (mTwf1) was expressed as a SNAP-tagged fusion protein and fluorescently labelled with bezylguanine-549, as done previously [27]. Photobleaching data confirmed that majority of 549-mTwf1 molecules were labeled with only one dye molecule, consistent with a monomeric nature of twinfilin [27]. Alexa-488 labelled actin filaments were polymerized in a non-microfluidic conventional open flow-cell and anchored on the glass coverslip using biotinylated-actin monomers. The filaments were then aged to ADP-F-actin, and subsequently exposed to fluorescently labeled mTwf1 (549-mTwf1). Twinfilin molecules preferentially associated with filament barbed ends rather than filament side (Supplementary Fig. 1). We observed repeated and transient short-lived appearances of 549-mTwf1 fluorescence signal at barbed ends (Fig. 1a). Notably, 549-mTwf1 intensity at the barbed end appeared and disappeared in a single step, indicating that only a single 549-mTwf1 molecule was present at a time at the barbed end (Fig. 1b,c). The barbed-end residence times of 549-mTwf1 molecules were exponentially distributed with an average dwell-time of about 0.54 ± 0.05 s (mean ± sem) (Fig. 1d).

**Fig. 1:**
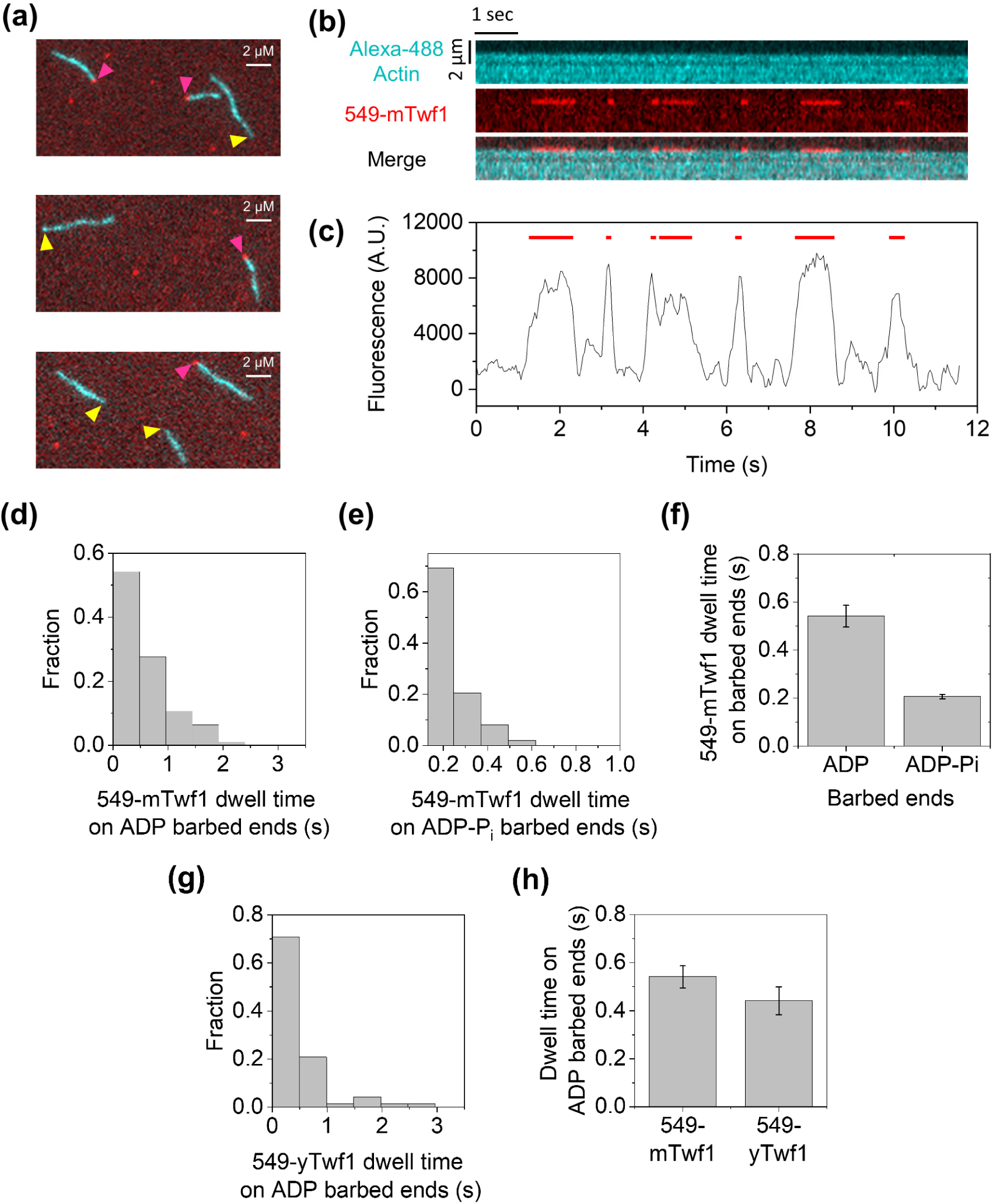
Direct visualization of mouse and yeast twinfilin at actin filament barbed ends. **(a)** Representative images of a multi-color single-molecule TIRF experiment. Actin filaments (green) were first assembled from 1 µM G-actin (15% Alexa-488 labeled, 0.5% biotin labeled) and 0.5 µM profilin. The chamber was washed with TIRF buffer, filaments were aged to ADP-F-actin state for 15 minutes and then subsequently exposed to 30 nM 549-mTwf1 (red). The yellow and magenta arrowheads indicate free and 549-mTwf1-bound barbed ends respectively. Scale bar, 2 µm. **(b)** Kymographs showing transient binding of 549-mTwf1 to filament barbed ends recorded at 0.041 s time resolution. Alexa-488 actin (cyan, top), 549-mTwf1 (red, middle) and merge (bottom). **(c)** Time record of 549-mTwf1 fluorescence intensity at the barbed end of a filament. Intensity is integrated over a 6 × 6 pixel square centered around the barbed end of the filament, and was smoothed using a moving window of 0.164 s (4 frames) **(d)** Distribution of barbed-end residence times of 549-mTwf1 on ADP-actin filaments (n= 94 binding events across 8 filaments). Mean residence time = 0.54 ± 0.04 s (± sem) **(e)** Distribution of barbed-end residence times of 549-mTwf1 on ADP-P_i_-actin filaments (n= 98 binding events across 7 filaments). Mean residence time = 0.20 ± 0.01 s (± sem) **(f)** Average residence times of 549-mTwf1 on ADP and ADP-P_i_ filament barbed ends **(g)** Distribution of residence times of 549-yTwf1 on ADP filament barbed ends (n= 72 binding events across 3 filaments). Mean dwell-time = 0.44 ± 0.06 s (± sem) (h) Average residence time of 549-mTwf1 and 549-yTwf1 on ADP filament barbed ends.

To our surprise, our measured mTwf1 barbed-end residence time was approximately two orders of magnitude lower than the previously reported values for *S. cerevisiae* twinfilin yTwf1 (∼69 s) [12]. To explore potential factors contributing to this discrepancy, we first considered the differences in the nucleotide state of actin filaments between the two studies. Notably, the Johnston et al study had not actively aged the actin filaments in their experiments. We wondered if this could have resulted in their filaments containing a mix of ADP and ADP-P_i_ actin subunits. We therefore repeated our single-molecule experiments using ADP-P_i_ actin filaments which were maintained in the ADP-P_i_ state by the addition of 50 mM P_i_, as done previously [13]. Remarkably, we observed a 2.5-fold reduction in the barbed-end residence time of twinfilin molecules (0.20 ± 0.01 s, mean ± sem) over ADP filaments (Fig. 1e,f). While our results are consistent with previous reports of higher affinity of twinfilin for ADP over ATP/ADP-P_i_ filaments [10], barbed-end residence time of mTwf1 still remained over two orders of magnitude shorter than the reported yTwf1 dwell-times.

We then considered whether the discrepancy might originate from species-specific differences which have previously been reported for twinfilin [28]. To test this, we purified and fluorescently labeled SNAP-yTwf1. Surprisingly, and contrary to previously reported long-lived interactions, our single-molecule experiments revealed that yeast twinfilin also exhibited transient interactions with filament barbed ends, displaying an average dwell-time of 0.44 s± 0.06 s (mean ± sem) (Fig.1 g,h). The underlying reasons for the discrepancy between our measured dwell times and those previously reported remain unclear. However, it is worth noting that in the Johnston et al. study, yeast twinfilin molecules were not freely diffusive but appear attached to the coverslip. This could have hindered the detection of twinfilin dissociation from barbed ends, potentially leading to re-association with barbed ends causing an appearance of longer-lived interactions. Another possibility is that the labeled twinfilin molecules in the study were actually oligomers, whose multivalent interactions with filament ends might have caused longer than usual dwell-times.

### Twinfilin is a non-processive barbed-end depolymerase

Twinfilin was previously suggested to be a processive depolymerase, meaning that each twinfilin molecule could continuously remove successive actin monomers while tracking a depolymerizing barbed end [12]. In light of our new data, we revisited this question by examining the number of actin subunits removed per binding event. Although Brownian motion-related fluctuations of twinfilin-bound barbed ends prevented a precise determination of filament length changes during individual binding events, we were able to calculate the average number actin subunits removed (from the barbed end) per mTwf1 binding event.

At saturating twinfilin concentrations, the filament barbed end is expected to be almost always occupied by a twinfilin molecule [13]. For ADP-P_i_ filaments, the maximum depolymerization rate observed at saturating twinfilin is ∼2.2 subunits per second at [13]. Given the average dwell-time of ∼0.2 s for 549-mTwf1 molecules at ADP-P_i_ barbed ends, approximately 0.4 actin subunits are lost from the barbed end during a single twinfilin binding event. This data suggests that not all twinfilin binding events at the barbed ends are productive, meaning every twinfilin molecule associating with the barbed end might not always cause removal of actin subunits. In case of ADP filaments, at saturating twinfilin concentration, barbed-ends depolymerized at 3.6 subunits per second [13]. Given the average dwell-time of 0.54 s for 549-mTwf1 molecules, about 1.9 actin subunits, on average, are lost per twinfilin binding event. Taken together, our analysis suggests that following their association with barbed ends, twinfilin molecules dissociate with either one or two actin subunits at a time. Thus, contrary to previous proposals, our data suggest that twinfilin is not a processive depolymerase i.e., twinfilin molecules do not track depolymerizing barbed ends.

### Twinfilin uncaps ADP filaments faster than ADP-P_i_ filaments

In light of our results demonstrating the actin filament age dependence on twinfilin’s barbed-end residence times, we investigated whether the uncapping ability of twinfilin also depended upon filament age. Previously, uncapping by twinfilin has only been investigated for ADP filaments [22]. To measure dissociation rate of CP, we employed an approach that has been previously used by us and others to measure dissociation rate of formin and CP from filament barbed ends [26, 27, 29, 30]. Pre-formed actin filaments were introduced into the mF-TIRF flow cell and captured by coverslip-anchored CP molecules (Fig. 2a). The filaments were then incubated with TIRF buffer (see methods) for 15 min to ensure their aging, before recording the time-dependent detachment of filaments from coverslip-bound CP and their subsequent disappearance from the field of view. The survival fraction of filaments was used to calculate the dissociation rate constant of CP from barbed ends. To study uncapping of ADP-P_i_ filaments, the filaments were maintained in a modified TIRF buffer, supplemented with 50 mM P_i_, throughout the experiment [13]. First, we sought to determine if the spontaneous off-rate of CP was influenced by filament age (in absence of twinfilin). By comparing the dissociation rates of CP from ADP and ADP-P_i_ barbed ends in the absence of twinfilin (see methods), we found that ADP filament barbed ends get uncapped ∼threefold faster than ADP-P_i_ filaments (Fig. 2c,d).

**Fig 2.**
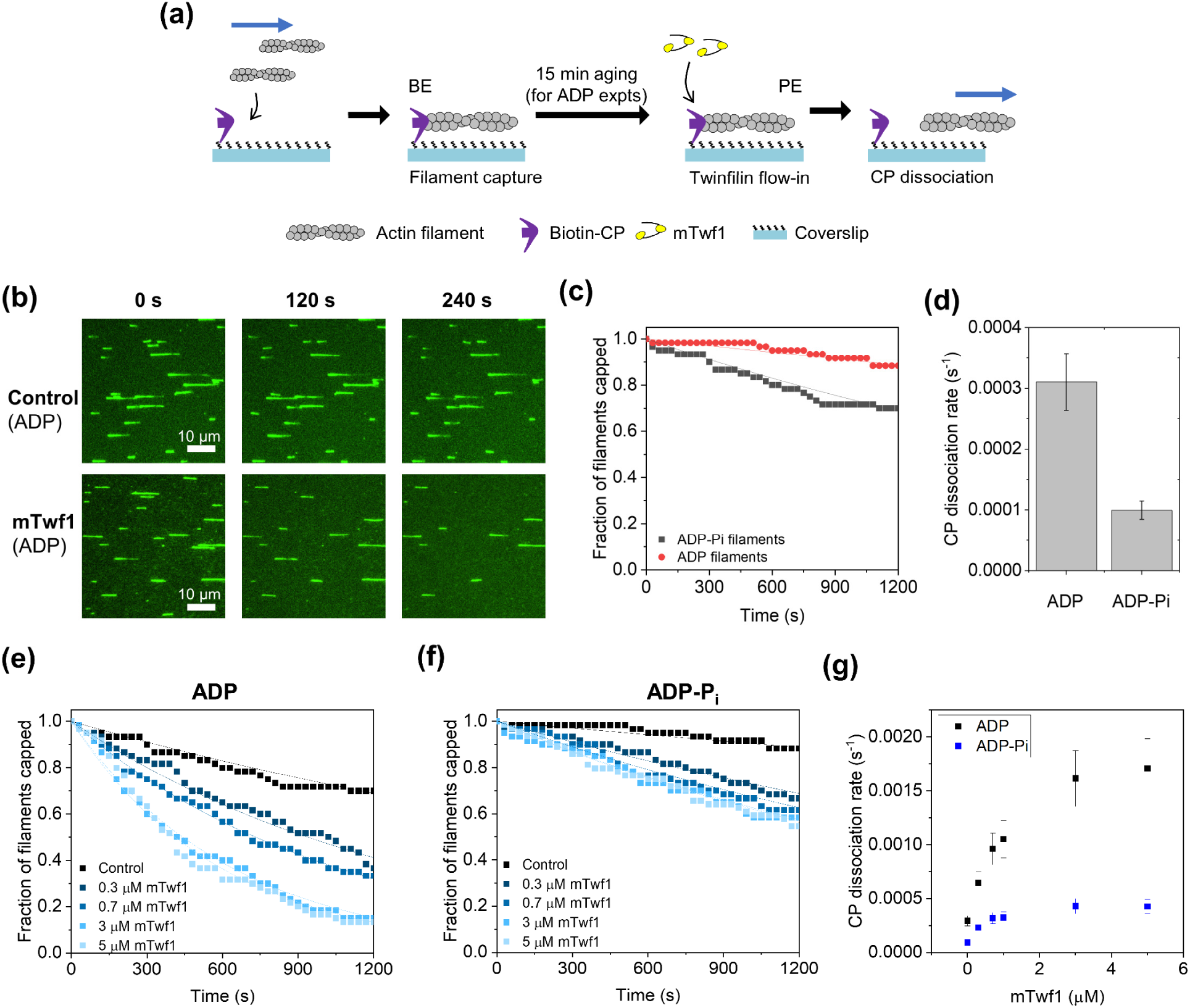
Twinfilin accelerates uncapping rate of both ADP and ADP-P_i_ filaments. **(a)** Schematic representation of the experimental strategy. Pre-formed Alexa-488 labeled actin filaments were captured by coverslip-anchored biotinylated SNAP-CP. The survival fraction of attached CP-bound filaments was monitored as a function of time in the presence of varying concentrations of mTwf1. For ADP-F-actin experiments, filaments were additionally aged for 15 min prior to their exposure to mTwf1. BE, barbed end; PE, pointed end. **(b)** Representative time-lapse images of a mF-TIRF field of view showing ADP-actin filaments (green) detaching from surface-anchored CP in presence of buffer (top) or 5 µM mTwf1 (bottom). Scale bar, 10 µm. **(c)** Survival fraction of CP-bound ADP and ADP-P_i_ actin filaments as a function of time in absence of mTwf1. **(d)** CP dissociation rate *k-_C_* for ADP and ADP-P_i_ actin filaments, determined from data in (c). **(e)** Survival fraction of CP-bound ADP actin filaments as a function of time in presence of a range of mTWf1 concentrations. **(f)** Survival fraction of CP-bound ADP-P_i_ actin filaments as a function of time in presence for a range of mTWf1 concentrations. In (c), (e) and (f), experimental data (symbols) are fitted to a single-exponential decay function (lines) to determine CP dissociation rate *k_-C_*. Number of filaments analyzed for each condition in (c), (e) and (f) : 60 **(g)** CP dissociation rate *k-_C_* for ADP and ADP-P_i_-actin filaments as a function of mTwf1 concentration, determined from data in (e) and (f).

When the reactions were supplemented with twinfilin, the uncapping rate of ADP filaments increased with twinfilin concentration until eventual saturation (Fig. 2b,g), consistent with previously published results [22]. Specifically, 5 µM mTwf1 increased the rate of CP dissociation from ADP barbed ends by about sixfold (Fig. 2e,g). To ensure that anchoring filaments with CP, or any flow-induced drag force didn’t influence the rate of uncapping, we compared the effects of twinfilin on filaments anchored at their pointed ends and capped at their free barbed end (Supplementary Fig. 2). Since the CP was now bound to the free distal ends of these filaments, these experiments mimicked previously-employed configuration by Hakala and colleagues [22]. These experiments yielded a sevenfold increase in uncapping by twinfilin, similar to the twinfilin-mediated enhancement observed in our CP-anchored experiments and aligning with previously published data [22]. Next, we investigated if twinfilin could also uncap ADP-P_i_ filaments. Similar to ADP filaments, the uncapping rate for ADP-P_i_ filaments increased with twinfilin concentration and saturated at high twinfilin concentrations. Compared to the control, 5 µM mTwf1 increased the rate of CP dissociation from ADP-P_i_ barbed ends by about fourfold (Fig. 2f,g). Furthermore, 5 µM twinfilin uncapped ADP filaments about fourfold faster than ADP-P_i_ filaments (Fig. 2g). Taken together, our analysis shows that capping protein stays bound longer to ADP-P_i_ filament ends than to ADP filament ends, both in the absence and presence of twinfilin.

### Formin uncaps ADP filaments faster than ADP-P_i_ filaments

We have previously demonstrated that, like twinfilin, formin can also displace CP from the barbed end [26, 27]. However, since neither of the previous studies controlled for filament age, the dependence of formin’s uncapping ability on filament age remains unclear. To address this, pre-formed actin filaments were captured by coverslip-bound CP and exposed to a solution with or without formin mDia1 (Fig. 3a). For ADP-F-actin experiments, the filaments were additionally aged for 15 min as described in the previous section. We systematically varied the concentration of formin and measured the barbed-end uncapping rates. Our results showed that for ADP filaments, the rates of barbed end uncapping increased linearly with formin concentration (Fig. 3c,e). At 200 nM formin, the highest concentration we tested, we observed a maximum uncapping rate of 0.01 s⁻¹, nearly a 35-fold increase over control reactions without formin (Fig. 3c,e).

**Fig 3.**
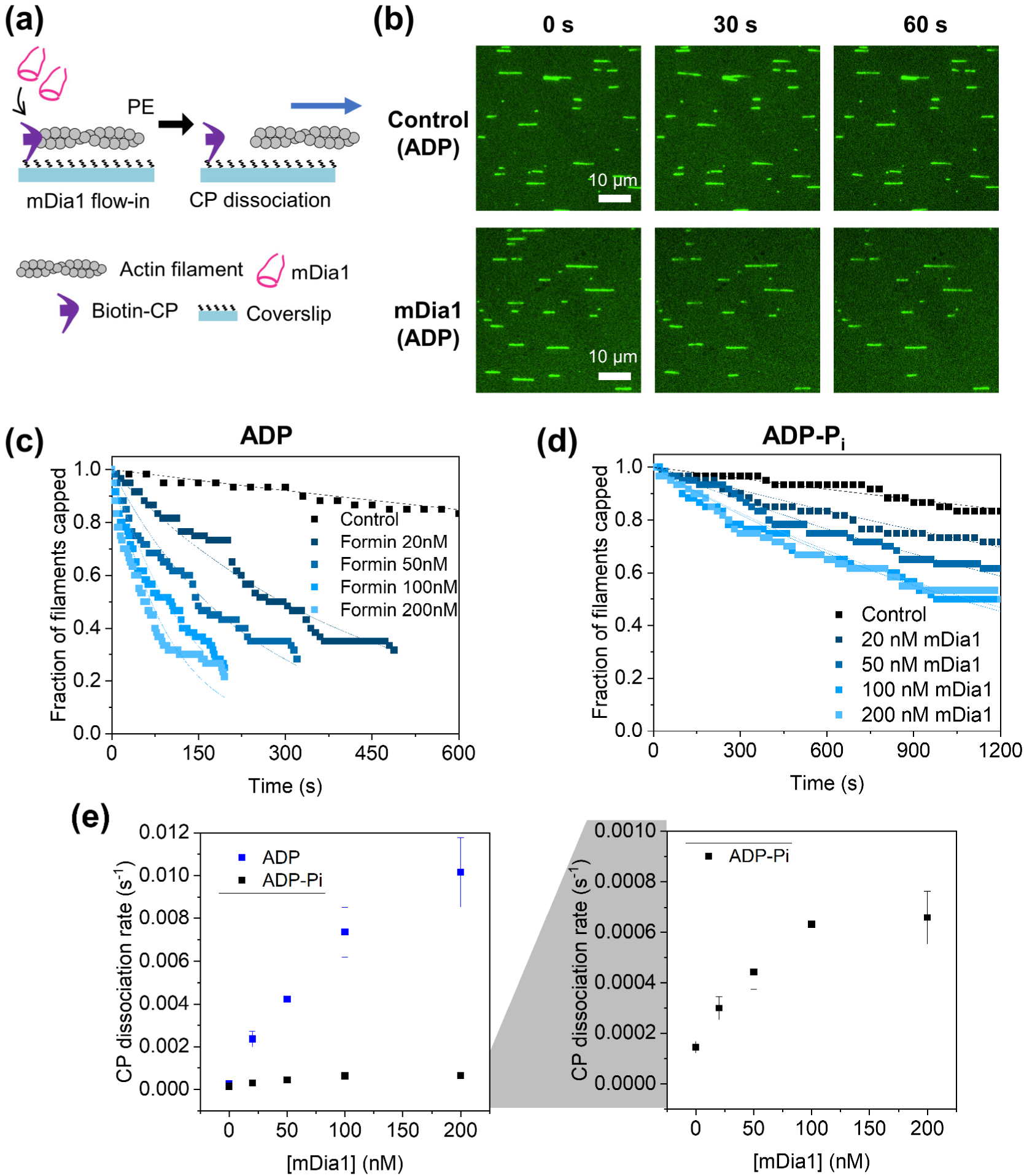
Formin accelerates uncapping rate of both ADP and ADP-P_i_ filaments. **(a)** Schematic representation of the experimental strategy, similar to shown in Fig. 2a. PE, pointed end. **(b)** Representative time-lapse images of an mF-TIRF field of view showing ADP-actin filaments (green) detaching from surface-anchored CP in presence of buffer (top) or 200 nM mDia1 (bottom). Scale bar, 10 µm. **(c)** Survival fraction of CP-bound ADP actin filaments as a function of time in presence for a range of mDia1 concentrations. **(d)** Survival fraction of CP-bound ADP-P_i_ actin filaments as a function of time in presence for a range of mDia1 concentrations. In (c) and (d), experimental data (symbols) are fitted to a single-exponential decay function (lines) to determine CP dissociation rate *k_-C_*. Number of filaments analyzed for each condition in (c) and (d): 60 **(e)** CP dissociation rate *k-_C_* for ADP and ADP-P_i_-actin filaments as a function of mDia1 concentration, determined from data in (c) and (d).

Interestingly, when we repeated this experiment with ADP-P_i_ filaments, we measured a maximum uncapping rate of 6.5 x 10^-4^ s^-1^, a sevenfold increase over buffer alone (Fig. 3d,e). Thus, although formin can accelerate uncapping independent of the nucleotide state of the actin filament, the absolute rate of uncapping differs between the two nucleotide states. Furthermore, the slower rate of uncapping for ADP-P_i_ filaments compared to ADP filaments follows the behavior of twinfilin-mediated uncapping. These results indicate that formin, like twinfilin, uncaps ADP filaments more efficiently than ADP-P_i_ filaments, highlighting the influence of the nucleotide state on the uncapping process.

### Twinfilin and formin synergize to accelerate uncapping of aged filaments but not ADP-P_i_ filaments

We recently demonstrated that twinfilin can join formin-CP “decision complex” at the barbed end. In the absence of twinfilin, both formin and CP departed with equal probabilities. However, the addition of twinfilin resulted in a higher fraction of decision complexes losing CP rather than formin [27]. We therefore decided to elucidate how uncapping rates might be influenced by simultaneous presence of twinfilin and formin, for both ADP and ADP-P_i_ filaments. Pre-formed actin filaments were captured at their barbed ends by coverslip-anchored CP and exposed to a solution containing both formin and twinfilin (Fig. 4a). We maintained twinfilin at a saturating concentration of 5 µM and systematically varied the formin concentration. For ADP filaments, the simultaneous presence of the two proteins drastically increased the uncapping rate compared to when either twinfilin or formin were present alone (Fig. 4b). Almost all the filaments in the field of view were uncapped in less than 60 s. Remarkably, twinfilin and formin together caused a ∼318-fold increase in the uncapping rate compared to the control reaction (buffer alone) (Fig. 4c,e,f). In presence of 5 µM twinfilin and 200 nM formin, we measured a 10-fold and 55-fold increase in uncapping rate over twinfilin alone and formin alone conditions (Fig. 4c,e,f).

**Fig 4.**
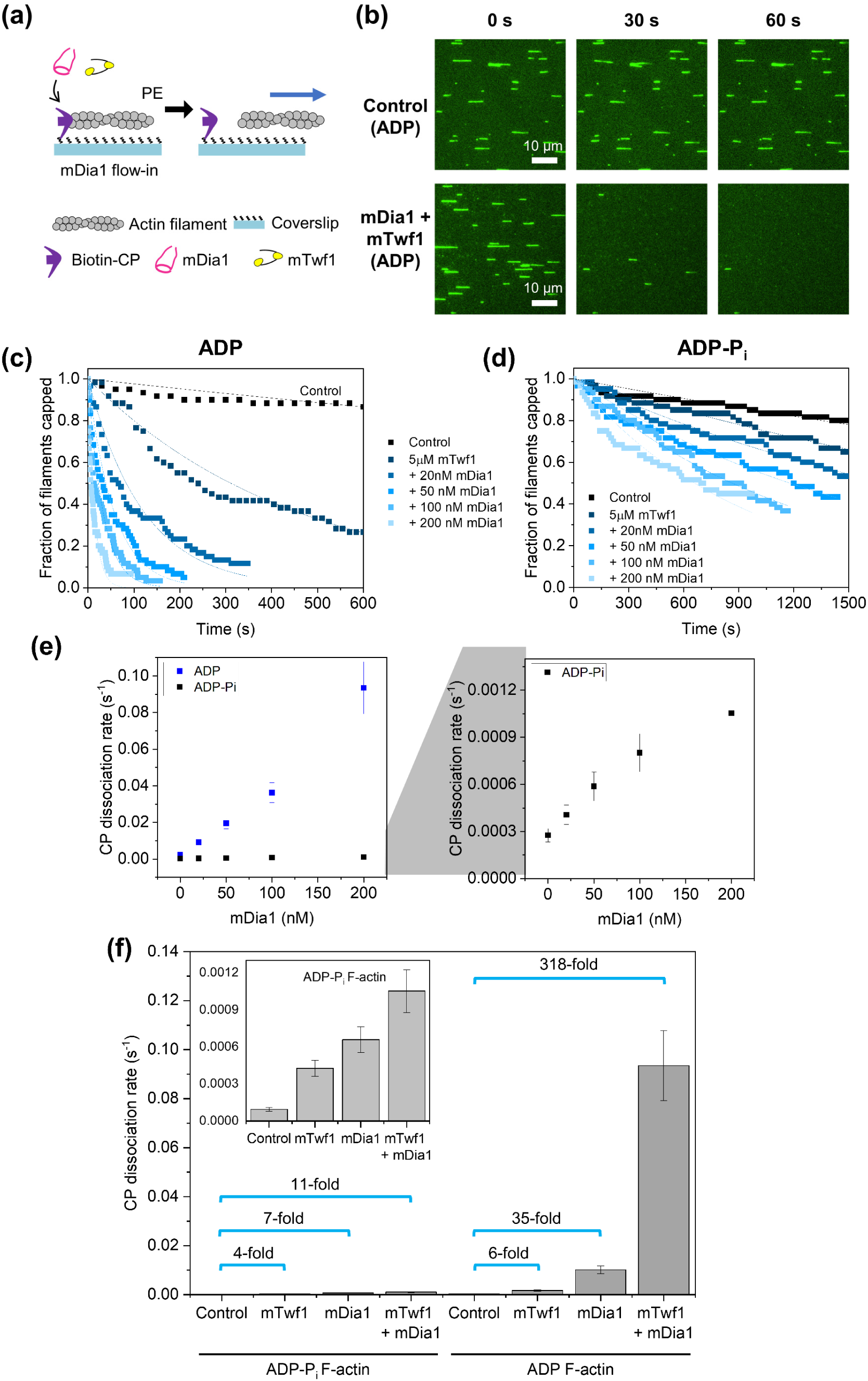
Formin and Twinfilin synergize to accelerate uncapping rate of both ADP and ADP-P_i_ filaments. **(a)** Schematic representation of the experimental strategy, similar to shown in Fig. 2a. PE, pointed end. **(b)** Representative time-lapse images of an mF-TIRF field of view showing ADP-actin filaments (green) detaching from surface-anchored CP in presence of buffer (top) or 200 nM mDia1 and 5 µM mTwf1 (bottom). **(c)** Survival fraction of CP-bound ADP actin filaments as a function of time in presence for a range of mDia1 concentrations (in presence of 5 µM mTwf1). **(d)** Survival fraction of CP-bound ADP-P_i_ actin filaments as a function of time in presence for a range of mDia1 concentrations (in presence of 5 µM mTwf1). In (c) and (d), experimental data (symbols) are fitted to a single-exponential decay function (lines) to determine CP dissociation rate *k_-C_*. Number of filaments analyzed for each condition in (c) and (d): 60. **(e)** CP dissociation rate *k-_C_* for ADP and ADP-P_i_-actin filaments as a function of mDia1 concentration (in presence of 5 µM mTwf1), determined from data in (c) and (d). **(f)** Side-by-side comparison of fold-changes in CP-dissociation rates by mDia1 and mTwf1 (alone and together) for ADP and ADP-Pi filaments (same data as shown in Figs. 1g, 2e and 3e).

Although the uncapping rates for ADP-P_i_ filaments also increased in the presence of both twinfilin and formin, the fold change was more modest compared to ADP filaments. At 5 µM twinfilin and 200 nM formin, we measured a 2.5-fold and 1.6-fold increase in uncapping rate over twinfilin alone or formin-alone conditions (Fig. 4d,e,f). Compared to buffer, the two proteins together caused an 11-fold enhancement in uncapping. These findings indicate that twinfilin and formin synergize to accelerate the uncapping of both ADP and ADP-P_i_ filament barbed ends, with a more pronounced effect on ADP filaments. Interestingly, when present together, the two proteins synergize to uncap ADP filaments, but their effect is only additive on ADP-P_i_ filaments. This multiprotein cooperation highlights the complex interplay between these proteins and their combined influence on actin filament dynamics.

## Discussion

Rapid actin depolymerization is essential for cells to dismantle older actin networks and regenerate monomers for new rounds of assembly[6]. Traditionally, actin filaments have been thought to depolymerize from their pointed ends by combined action of cofilin and its cofactor CAP [31–36]. While early studies identified twinfilin as G-actin sequestering protein and a barbed-end capper of actin filaments [7, 10, 11], a more recent single-molecule study instead suggested that twinfilin primarily acts as a processive depolymerase at filament barbed ends [12]. Later studies, however, have questioned twinfilin’s ability to processively depolymerize barbed ends, as discussed below.

First, insights from a recent X-ray structure of the actin-twinfilin-capping protein complex suggested that the twinfilin-CP complex may act as a transient capper, dissociating from the barbed end along with the terminal actin subunits [14]. Secondly, our previous observations showed that increasing amounts of G-actin (in the presence of saturating twinfilin) can modulate the rate of twinfilin-mediated depolymerization, suggesting that twinfilin and G-actin compete for barbed-end binding [13]. Consequently, barbed ends are expected to rapidly alternate between twinfilin-bound and bare states, with the fraction of time spent in each state determining the net elongation rate. Thirdly, a recent study which measured barbed-end depolymerization rates due to combined action of twinfilin, profilin and cofilin, found that while profilin and cofilin can simultaneously occupy a barbed end, twinfilin and profilin bind barbed ends mutually exclusively [37]. Based on these observations, the authors suggested that at the barbed end, twinfilin interacts with both the terminal actin subunits. In light of these conflicting findings, the question of whether twinfilin acts as a processive depolymerase remains unresolved.

Our study finds that both yeast and mouse twinfilin display short-lived associations with filament barbed ends, sensitive to filament age (Fig. 1b, d, g). Mouse twinfilin remains bound more tightly to ADP barbed ends (*t_dwell_* ∼0.54 seconds) than to ADP-P_i_ barbed ends (t_dwell_∼0.20 seconds). These observations are consistent with a previous study which reported a higher affinity of twinfilin for filament barbed ends under depolymerization conditions (K_d_ = 13 nM) than under assembly-promoting conditions (K_d_ = 0.1–0.3 µM), suggesting a preference of ADP filaments over ATP- or ADP-P_i_ filaments [10]. Other ADF-H family members, such as cofilin and GMF, also display a higher affinity for ADP-actin and ADP-Arp2/3 branches respectively [38–40].

Our analysis suggests that most twinfilin binding events result in removal of one or two actin subunits, with many binding events (especially on ADP-P_i_ filaments) being unproductive. This resembles our previous observations where multiple (∼30) short-lived twinfilin binding events were required to displace CP from barbed ends [27]. Twinfilin’s dwell-time on free barbed ends is ∼4 – 10 fold shorter than its dwell-time on CP-bound ends (∼1.9 s) [27], suggesting dual interactions of twinfilin with CP and terminal actin subunits stabilize its localization at filament barbed ends, leading to longer dwell-times on CP-bound ends. Our observations are consistent with similar suggestions by Mwangangi and colleagues [14].

Combining our results with previous observations, we propose a working model for twinfilin’s action at barbed ends (Fig. 5a,b). Twinfilin binds free barbed ends, acting as both a transient capper and depolymerase. In its capping role, twinfilin prevents filament elongation, while its depolymerase function removes terminal actin subunits. Without G-actin, twinfilin acts solely as a depolymerase. In the presence of high G-actin concentration, the barbed end switches between elongation, capping, and depolymerization states, depending on twinfilin’s presence or absence at the barbed end. Our findings thus resolve twinfilin’s long-debated barbed-end depolymerization mechanism.

**Fig. 5.**
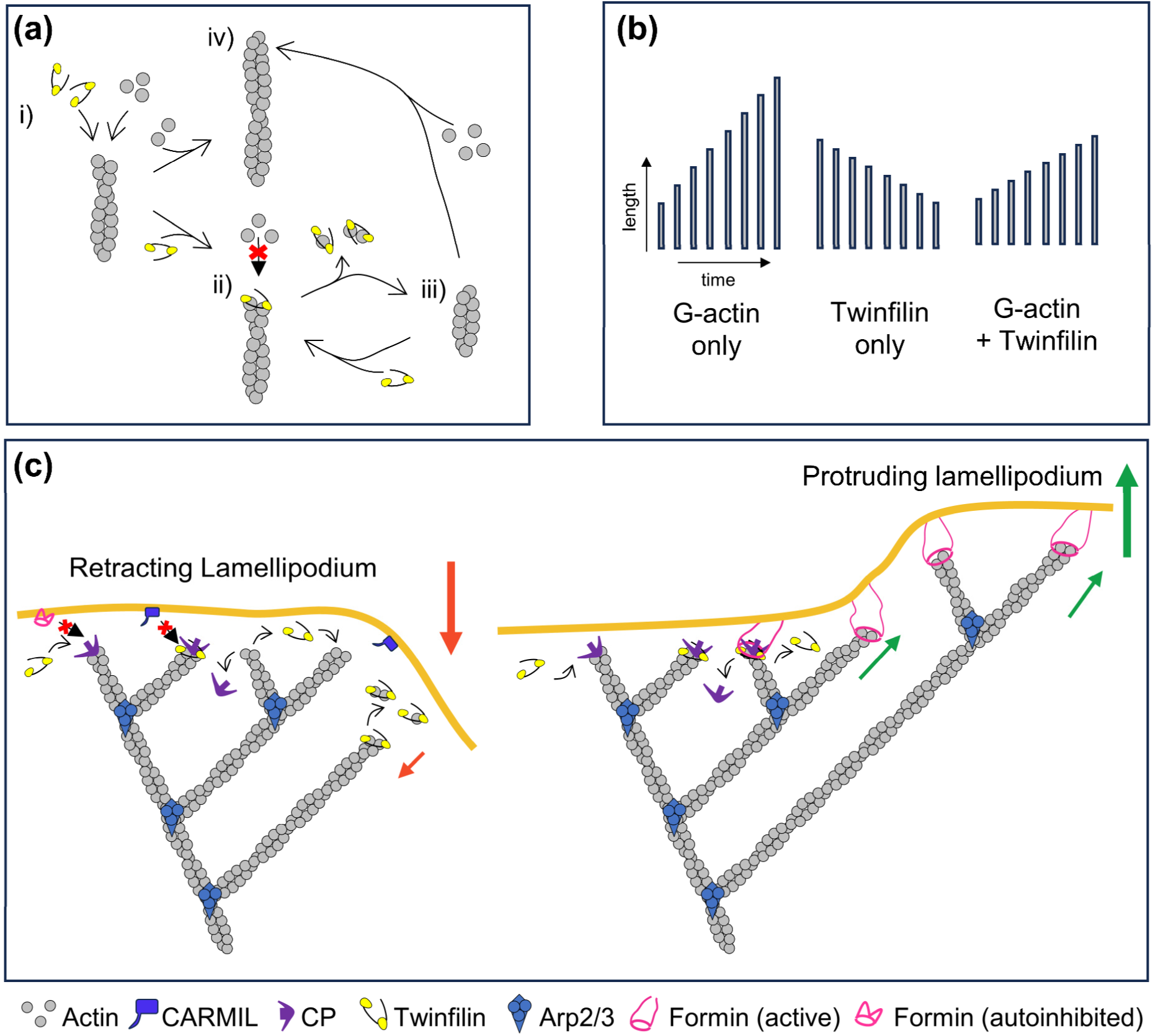
Working model for twinfilin’s multifaceted roles in filament capping, depolymerization and uncapping. **(a)** Effect of twinfilin on barbed-end dynamics in simultaneous presence of actin monomers and twinfilin : (i) If a twinfilin molecule binds to the barbed end, it acts as a transient capper and actin monomers cannot add to the twinfilin occupied barbed end (ii). Twinfilin then leaves with either one or two actin monomers. Following twinfilin’s departure, either another twinfilin molecule could occupy the barbed end resulting in depolymerization (iii) or actin monomers could bind to the free barbed end resulting in polymerization (iv). **(b)** Schematic representation of time-dependent changes in filament length in presence G-actin and twinfilin (alone or together). The filaments elongate fast if only actin monomers are present (left), depolymerize if only twinfilin is present (middle), and grow at a slower rate if both twinfilin and actin monomers are present and compete for the barbed end (right). **(c)** CP, formin, twinfilin, and CARMIL all localize to the leading edge of a motile cell. CP-bound barbed ends could have two different fates depending on whether they get exposed to twinfilin or twinfilin and formin. In a retracting lamellipodium (left), twinfilin only modestly uncaps the CP-capped barbed ends, it protect CP from CARMIL – thus preventing rapid uncapping and filament regrowth. The uncapped barbed ends can then get depolymerized with the help of twinfilin which will lead to lamellipodium retraction. However, when both formin and twinfilin are present, they can rapidly uncap CP-capped barbed end. CP and twinfilin depart from the end, leaving formin behind which can reinitiate rapid elongation which will result in switching a retracting lamellipodium into a protruding one (right). Red and green arrows represent retraction and protrusion respectively.

Twinfilin also impacts cellular localization and dynamics of CP [22]. It binds CP via its CPI motif and accelerates CP dissociation sixfold[20–22]. However, these observations do not fully explain the much faster dissociation of CP observed in cells, (*k_off_* = 0.14 – 0.58 s^-1^) [18, 19] suggesting additional factors work with twinfilin in vivo. We find that twinfilin synergizes with formin to accelerate uncapping by up to 318-fold, with rates depending on filament age. Our study is the first to link filament age with CP dynamics at the filament end, highlighting the importance of nucleotide state in actin filament turnover.

Where in a cell might these mechanisms be relevant? These mechanisms are likely crucial in regions requiring rapid actin reorganization, like the leading edge of migrating cells where CP, twinfilin, formin and CARMIL all localize [18, 22, 41, 42](Fig. 5c). Lamellipodia undergo cycles of leading-edge protrusion and retraction. During retraction, actin filament growth needs to be arrested and elongators like formin are expected to be auto-inhibited. In such a situation, twinfilin would protect CP from being uncapped by CARMIL, preventing filament regrowth. In cases when a retracting lamellipodia needs to rapidly switch to protruding, active formin and twinfilin can synergize to rapidly uncap CP-capped filaments and reinitiate their elongation. Our results also indicate that ADP filament barbed ends display significantly faster uncapping rates compared to ADP-P_i_ filaments. Although P_i_ release is slow in the bulk of the filaments, it occurs 240-fold faster at the barbed-end subunits (t_1/2_ = 0.39 s) [43]. Consistently, Funk et al. observed a lack of P_i_ density at terminal barbed-end subunits of CP-capped filaments in their cryo-EM study [44]. Interestingly, our observations suggest that binding of just a single twinfilin molecule at the growing barbed end (t_dwell_ ∼0.2 – 0.54 s) (resulting in arrest of growth during that period) might provide sufficient time to facilitate conversion of terminal actin subunits into ADP-actin. Together, these findings suggest that just as at free barbed ends, the terminal and penultimate subunits might also rapidly experience P_i_ release on capped barbed ends and get quickly converted into ADP-actin subunits. This could potentially lead to faster uncapping of recently-assembled capped filaments by mechanisms uncovered here. In summary, our findings provide insights into twinfilin’s multifunctional barbed-end roles as a depolymerase, transient capper, and uncapper which synergizes with formin to rapidly displace CP.

## Methods

### Purification and labeling of actin

Rabbit skeletal muscle actin was purified from acetone powder generated from frozen ground muscle tissue of young rabbits. Lyophilized acetone powder stored at −80°C was mechanically sheared in a coffee grinder, resuspended in G-buffer (5 mM Tris-HCl pH 7.5, 0.5 mM Dithiothreitol (DTT), 0.2 mM ATP and 0.1 mM CaCl_2_), and cleared by centrifugation for 20 min at 50,000 × *g*. Supernatant was collected and further filtered with Whatman paper. Actin was then polymerized overnight at 4°C, slowly stirring, by the addition of 2 mM MgCl_2_ and 50 mM NaCl to the filtrate. The next morning, NaCl powder was added to a final concentration of 0.6 M and stirring was continued for another 30 min at 4°C. Then, F-actin was pelleted by centrifugation for 150 min at 280,000 × *g*, the pellet was solubilized by dounce homogenization and dialyzed against G-buffer for 48 h at 4°C. Monomeric actin was then precleared at 435,000 × *g* and loaded onto a Sephacryl S-200 16/60 gel-filtration column (Cytiva, USA) equilibrated in G-Buffer. Fractions containing actin were stored at 4°C.

To fluorescently label actin, G-actin was polymerized by dialyzing overnight against modified F-buffer (20 mM PIPES pH 6.9, 0.2 mM CaCl_2,_ 0.2 mM ATP, 100 mM KCl) [45]. F-actin was incubated for 2 h at room temperature with a 5-fold molar excess of Alexa-488 NHS ester dye (Thermo Fisher Scientific, USA). F-actin was then pelleted by centrifugation at 450,000 × *g* for 40 min at room temperature, and the pellet was resuspended in G-buffer, homogenized with a dounce and incubated on ice for 2 h to depolymerize the filaments. The monomeric actin was then re-polymerized on ice for 1 h by addition of 100 mM KCl and 1 mM MgCl_2_. F-actin was once again pelleted by centrifugation for 40 min at 450,000 × *g* at 4°C. The pellet was homogenized with a dounce and dialyzed overnight at 4°C against 1 L of G-buffer. The solution was precleared by centrifugation at 450,000 × *g* for 40 min at 4°C. The supernatant was collected, and the concentration and labeling efficiency of actin was determined.

### Purification and labeling of mTwf1 and yTwf1

Mouse twinfilin (mTwf1) was expressed in *E. coli* BL21 (pRare). Cells were grown in Terrific Broth to log phase at 37°C. Expression was induced overnight at 18°C by addition of 1 mM IPTG. Cells were harvested by centrifugation at 11,200 × *g* for 15 min and the cell pellets were stored at - 80°C. For purification, frozen pellets were thawed and resuspended in 35 mL lysis buffer (50 mM sodium phosphate buffer pH 8, 20 mM imidazole, 300 mM NaCl, 1 mM DTT, 1 mM PMSF and protease inhibitors (pepstatin A, antipain, leupeptin, aprotinin, and chymostatin, 0.5 μM each)). Cells were lysed using a tip sonicator while kept on ice. The cell lysate was then centrifuged at 120,000 × *g* for 45 min at 4°C. The supernatant was then incubated with 1 mL of Ni-NTA beads (Qiagen, USA) while rotating for 2 h at 4°C. The beads were then washed three times with the wash buffer (50 mM sodium phosphate buffer pH 8, 300 mM NaCl, 20 mM imidazole and 1 mM DTT). The beads were then transferred to a disposable column (Bio-Rad, USA). Protein was eluted using the elution buffer (50 mM phosphate buffer pH 8, 300 mM NaCl, 250 mM imidazole and 1 mM DTT). Fractions containing the protein were concentrated and loaded onto a size exclusion Superdex 75 Increase 10/300 column (Cytiva, USA) pre-equilibrated with 20 mM HEPES pH 7.5, 1 mM EDTA, 50 mM KCl and 1 mM DTT. Peak fractions were collected, concentrated, aliquoted, and flash-frozen in liquid N_2_ and stored at −80°C.

SNAP-mTwf1 [27] and SNAP-yTwf1 [12] was purified using the same protocol as above. Purified SNAP-mTwf1 and SNAP-yTwf1 were incubated with 5x excess of SNAP-surface-549 dye (New England Biolabs, Ipswich, MA) overnight at 4°C. Free dye was removed using a PD-10 desalting column (Cytiva, USA). Labeled protein was collected, concentrated, aliquoted, and flash-frozen in liquid N_2_ and stored at −80°C.

### Purification of formin mDia1

Mouse his-tagged mDia1 (FH1-FH2-C) formin was expressed in *E. coli*; BL21(DE3) pLysS cells. Cells were grown in Terrific Broth to log phase at 37°C. Expression was induced overnight at 18°C by addition of 1 mM IPTG. Cells were harvested by centrifugation at 11,200 × *g* for 15 min and the cell pellets were stored at −80°C. For purification, frozen pellets were thawed and resuspended in 35 mL lysis buffer (50 mM sodium phosphate buffer pH 8, 20 mM imidazole, 300 mM NaCl, 1 mM DTT, 1 mM PMSF and protease inhibitors (0.5 μM each of pepstatin A, antipain, leupeptin, aprotinin, and chymostatin)). Cells were lysed using a tip sonicator while being kept on ice. The cell lysate was then centrifuged at 120,000 × *g* for 45 min at 4°C. The supernatant was then incubated with 1 mL of Ni-NTA beads (Qiagen, USA) while rotating for 2 h at 4°C. The beads were then washed three times with the wash buffer (50 mM sodium phosphate buffer pH 8, 300 mM NaCl, 20 mM imidazole and 1 mM DTT) and were then transferred to a disposable column (Bio-Rad, USA). Protein was eluted using the elution buffer (50 mM phosphate buffer pH 8, 300 mM NaCl, 250 mM imidazole and 1 mM DTT). Fractions containing the protein were concentrated and loaded onto a size exclusion Superdex 200 increase 10/300 GL column (Cytiva, USA) pre-equilibrated with 20 mM HEPES pH 7.5, 150 mM KCl, 10% glycerol and 0.5 mM DTT. Peak fractions were collected, concentrated, aliquoted, and flash-frozen in liquid N_2_ and stored at - 80°C.

### Purification and biotinylation of capping protein

Mouse his-tagged capping protein was expressed in *E. coli*; BL21(DE3) pLysS cells. Capping protein subunits α1 and β2 were expressed from the same plasmid with a single His-tag on the alpha subunit [26]. Cells were grown in Terrific Broth to log phase at 37°C. Expression was induced overnight at 18°C by addition of 1 mM IPTG. Cells were harvested by centrifugation at 11, 200 × *g* for 15 min and the cell pellets were stored at −80°C. For purification, frozen pellets were thawed and resuspended in 35 mL lysis buffer (50 mM sodium phosphate buffer pH 8, 20 mM imidazole, 300 mM NaCl, 1 mM DTT, 1 mM PMSF and protease inhibitors (0.5 μM each of pepstatin A, antipain, leupeptin, aprotinin, and chymostatin)). Cells were lysed using a tip sonicator while being kept on ice. The cell lysate was then centrifuged at 120,000 × *g* for 45 min at 4°C. The supernatant was incubated with 1 mL of Ni-NTA beads (Qiagen, USA) while rotating for 2 h at 4°C. The beads were then washed three times with the wash buffer (50 mM sodium phosphate buffer pH 8, 300 mM NaCl, 20 mM imidazole and 1 mM DTT) and transferred to a disposable column (Bio-Rad, USA). Protein was eluted using an elution buffer (50 mM phosphate buffer pH 8, 300 mM NaCl, 250 mM Imidazole and 1 mM DTT). Fractions containing the protein were concentrated and loaded onto a size exclusion Superdex75 Increase 10/300 column (Cytiva, USA) pre-equilibrated with 20 mM Tris-HCl, 50 mM KCl and 1 mM DTT. Peak fractions were collected, concentrated, aliquoted, and flash-frozen in liquid N_2_ and stored at −80°C.

SNAP-CP was expressed from a single plasmid containing His- and SNAP-tagged β1 subunit and untagged α1 subunit [25]. It was purified using the protocol above. Purified SNAP-CP was incubated with 5x excess of SNAP-surface-Biotin (New England Biolabs, USA) overnight at 4°C. Free biotin were removed using PD-10 desalting columns (Cytiva, USA). Labeled protein was collected, concentrated, aliquoted, and flash-frozen in liquid N_2_ and stored at −80°C.

### Purification of profilin

Human profilin-1 was expressed in *E. coli* strain BL21 (pRare) to log phase in LB broth at 37°C and induced with 1 mM IPTG for 3 h at 37°C. Cells were then harvested by centrifugation at 15,000 × *g* at 4°C and stored at −80°C. For purification, pellets were thawed and resuspended in 30 mL lysis buffer (50 mM Tris-HCl pH 8, 1 mM DTT, 1 mM PMSF protease inhibitors (0.5 μM each of pepstatin A, antipain, leupeptin, aprotinin, and chymostatin)) was added, and the solution was sonicated on ice by a tip sonicator. The lysate was centrifuged for 45 min at 120,000 × *g* at 4°C. The supernatant was then passed over 20 ml of Poly-L-proline conjugated beads in a disposable column (Bio-Rad, USA). The beads were first washed at room temperature in wash buffer (10 mM Tris pH 8, 150 mM NaCl, 1 mM EDTA and 1 mM DTT) and then washed again with 2 column volumes of 10 mM Tris pH 8, 150 mM NaCl, 1 mM EDTA, 1 mM DTT and 3 M urea. Protein was then eluted with 5 column volumes of 10 mM Tris pH 8, 150 mM NaCl, 1 mM EDTA, 1 mM DTT and 8 M urea. Pooled and concentrated fractions were then dialyzed in 4 L of 2 mM Tris pH 8, 0.2 mM EGTA, 1 mM DTT, and 0.01% NaN_3_ (dialysis buffer) for 4 h at 4°C. The dialysis buffer was replaced with fresh 4 L buffer and the dialysis was continued overnight at 4°C. The protein was centrifuged for 45 min at 450,000 × *g* at 4°C, concentrated, aliquoted, flash frozen in liquid N_2_ and stored at −80°C.

### Conventional TIRF microscopy for single molecule imaging

Glass coverslips (60 x 24 mm; Thermo Fisher Scientific, USA) were first cleaned by sonication in detergent for 20 min, followed by successive sonications in 1 M KOH, 1 M HCl and ethanol for 20 min each. Coverslips were then washed extensively with H_2_O and dried in an N_2_ stream. The cleaned coverslips were coated with 2 mg/mL methoxy-polyethylene glycol (mPEG)-silane MW 2,000 and 2 µg/mL biotin-PEG-silane MW 3,400 (Laysan Bio, USA) in 80% ethanol (pH 2.0) and incubated overnight at 70°C. Flow cells were assembled by rinsing PEG-coated coverslips with water, drying with N_2_, and adhering to μ-Slide VI0.1 (0.1 mm x 17 mm x 1 mm) flow chambers (Ibidi, Germany) with double-sided tape (2.5 cm x 2 mm x 120 μm) and epoxy resin for 5 min (Devcon, USA). Before each reaction, the flow cell was sequentially incubated for 1 min each with 4 μg/ml streptavidin and 1% BSA in 20 mM HEPES pH 7.5, and 50 mM KCl. The flow cell was then equilibrated with TIRF buffer (10 mM imidazole, pH 7.4, 50 mM KCl, 1 mM MgCl_2_, 1 mM EGTA, 0.2 mM ATP, 10 mM DTT, 2 mM DABCO and 0.5% methylcellulose [4,000 cP]).

For two color experiments with 549-mTwf1 and 488-actin (Fig. XX), actin filaments were elongated from 1 µM G-actin (15% Alexa-488 labeled and 1.4% biotinylated G-actin) and 0.5 µM Profilin. Free actin monomers were removed by rinsing the flow cell with excess of TIRF buffer. For ADP filaments, filaments were aged by incubating them in TIRF buffer containing 0.1 µM G-actin for 15 minutes prior to being rinsed with TIRF buffer and introduction of 30 nM 549-mTwf1 or 549-yTwf1. Time-lapse images were acquired every 0.041 second. For ADP-P_i_ filaments, modified TIRF buffer (regular TIRF buffer supplemented with 50 mM inorganic phosphate : 10 mM imidazole pH 7.4, 34.8 mM K_2_HPO_4_ and 15.2 mM KH_2_PO_4_, 1 mM MgCl_2_, 1 mM EGTA, 0.2 mM ATP, 10 mM DTT, 1 mM DABCO) was used. The presence of 50 mM P_i_ in the TIRF buffer ensures that filaments remain in ADP-P_i_ state throughout the experiment.

### Microfluidics-assisted TIRF (mf-TIRF) microscopy

Uncapping of actin filaments was investigated using microfluidics-assisted TIRF (mf-TIRF) microscopy [45]. For all experiments, coverslips were first cleaned by sonication in Micro90 detergent for 20 min, followed by successive 20 min sonications in 1 M KOH, 1 M HCl and 200 proof ethanol for 20 min each. Washed coverslips were then stored in fresh 200 proof ethanol. Coverslips were then washed extensively with H_2_O and dried in an N_2_ stream. These dried coverslips were coated with 2 mg/mL methoxy-poly (ethylene glycol) (mPEG)-silane MW 2,000 and 2 µg/mL biotin-PEG-silane MW 3,400 (Laysan Bio, USA) in 80% ethanol (pH 2.0) and incubated overnight at 70°C. A 40 µm high PDMS mold with 3 inlets and 1 outlet was mechanically clamped onto a PEG-Silane coated coverslip. The chamber was then connected to a Maesflo microfluidic flow-control system (Fluigent, France), rinsed with TIRF buffer (10 mM imidazole pH 7.4, 50 mM KCl, 1 mM MgCl_2_, 1 mM EGTA, 0.2 mM ATP, 10 mM DTT, 1 mM DABCO) and incubated with 1% BSA and 10 µg/mL streptavidin in 20 mM HEPES pH 7.5, and 50 mM KCl for 5 min. Biotin-CP was first attached on the glass coverslip by flowing in 10 nM biotin-CP for 30 s. Pre-formed actin filaments (1 µM 15% Alexa-488 labelled) were then introduced and captured by anchored CP at their barbed ends with their distal pointed ends free in solution. For ADP experiments, actin filaments were first aged under continuous flow for 15 min and then exposed to specific biochemical conditions. For ADP-P_i_ filament experiments, modified TIRF buffer (regular TIRF buffer supplemented with 50 mM inorganic phosphate : 10 mM imidazole pH 7.4, 34.8 mM K_2_HPO_4_ and 15.2 mM KH_2_PO_4_, 1 mM MgCl_2_, 1 mM EGTA, 0.2 mM ATP, 10 mM DTT, 1 mM DABCO) was used. Presence of 50 mM P_i_ in the TIRF buffer ensures that filaments remain in ADP-P_i_ state throughout the experiment [4, 5]. For all experiments, time-dependent detachment of filaments from coverslip-bound CP was recorded. The survival fraction of filaments was used to calculate the dissociation rate constant of CP from barbed ends. All experiments were conducted at room temperature and under continuous flow which was maintained throughout the experiment.

### Image acquisition and analysis

Single-wavelength time-lapse TIRF imaging was performed on a Nikon-Ti2000 inverted microscope equipped with a 40 mW 488 nm Argon laser, a 60X TIRF-objective with a numerical aperture of 1.49 (Nikon Instruments Inc., USA) and an IXON LIFE 888 EMCCD camera (Andor Ixon, UK). One pixel was equivalent to 144 × 144 nm. Focus was maintained by the Perfect Focus system (Nikon Instruments Inc., Japan). Time-lapsed images were acquired using Nikon Elements imaging software (Nikon Instruments Inc., Japan). For two color images, the samples were simultaneously excited by 488 nm and 561 nm lasers and the emission was imaged on a single camera using an OptoSplit (Cairn Research). Time lapse images were acquired every 0.041s. A 6 x 6 pixel box was drawn at the barbed end of the filament and the time-dependent intensity values were recorded for the 549 nm channel. Only binding events that lasted longer than 3 frames were counted for calculation of average residence times. As a result, the calculated dwell-times are potentially being overestimated.

Images were analyzed in Fiji [46]. A kymograph plugin was used to make kymographs of individual filaments. These kymographs were then used to identify the timepoint of detachment. For each condition, 60 filaments were analyzed across multiple fields of view. All of these filaments were included to determine the cumulative distribution functions (CDFs) showing time dependent survival fraction of capped filaments. Data analysis and curve fitting were carried out in Microcal Origin. All experiments were repeated three at least times and yielded similar results. Data shown are from one trial.

### Statistical analysis and error bars for dissociation rates

The uncertainty in dissociation rates of CP were determined by bootstrapping strategy [27, 29]. The dissociation rate was determined by fitting the survival fraction (or CDF) data to a single exponential function (y = e^-*k*t^ or y = 1 – e^-*k*t^). A custom-written MATLAB code was then used to simulate CP lifetimes at barbed ends for N filaments (where N is the number of filaments in the particular experiment) based on the rate k determined from the experimental data. The simulation was repeated 1000 times to generate 1000 individual survival fractions of N filaments. Each dataset was then fit to an exponential function and an observed rate constant *k_obs_* was determined for each of the 1000 simulated datasets. The standard deviation of these estimated rates allowed us to determine the uncertainty in our measured rates.

## Data availability

Data supporting the findings of this manuscript are available from the corresponding author upon reasonable request.

## Code availability

Code used in this manuscript is available from the corresponding author upon reasonable request.

## Acknowledgements

This work was supported by NIH NIGMS grant R35GM143050 to SS.

## Author contributions

VR and AA conducted experiments. SS designed experiments and supervised the project. All authors analyzed data, prepared figures and wrote the manuscript. SS acquired funding.

---

**Supplementary Fig. 1:**
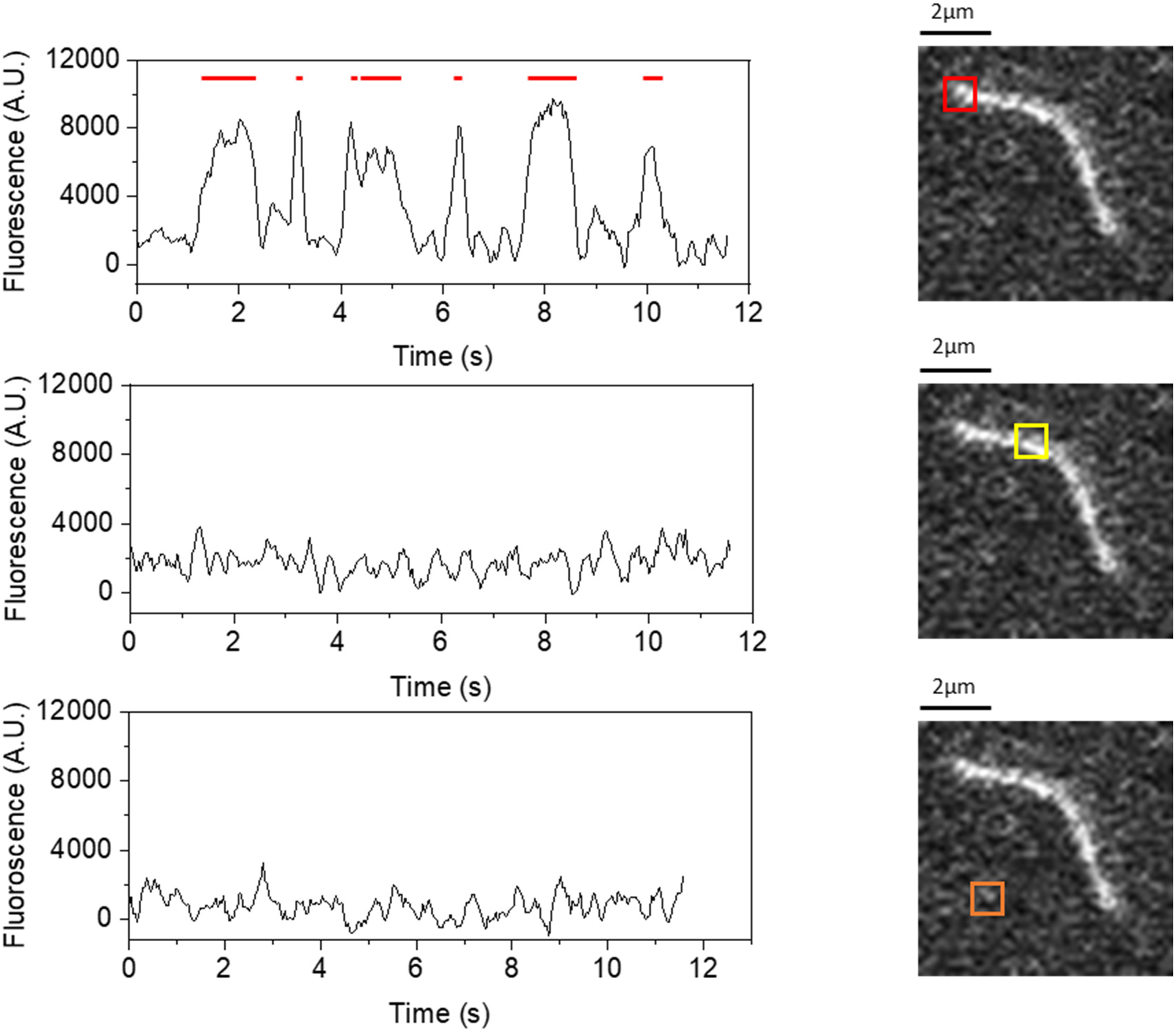
549-mTwf1 molecules preferentially bind to the barbed ends of actin filaments rather than to their sides. Time records of integrated 549-mTwf1 intensity from 6 x 6 pixel squares placed the following three regions – at the filament barbed end (top), a control region on the filament but away from the barbed end (middle), and on the coverslip in a region away from the filament (bottom). Intensity records shown were acquired at 0.041 s per frame and smoothed using an adjacent averaging algorithm over a 0.164 s sliding window.

**Supplementary Fig. 2:**
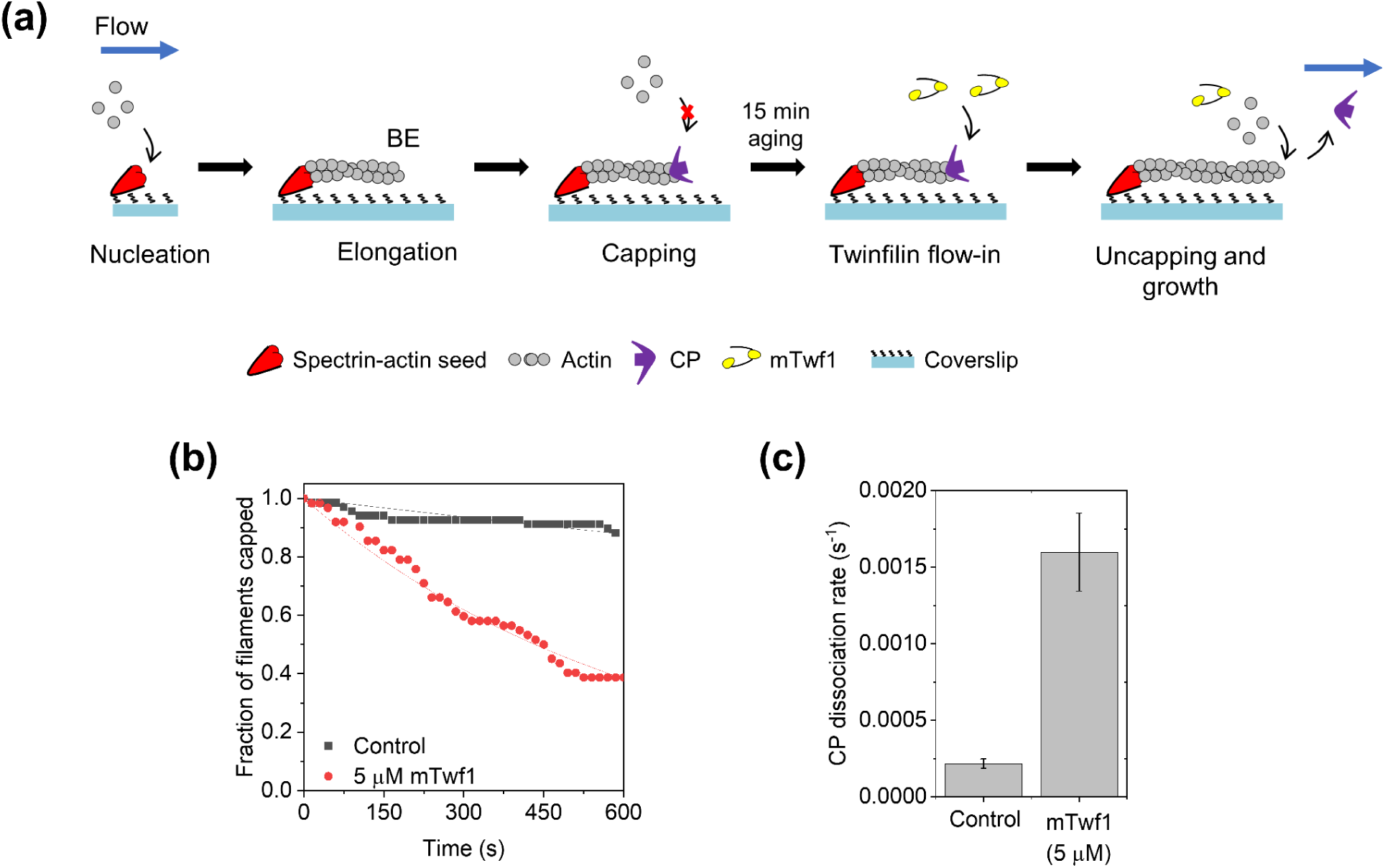
Uncapping by twinfilin of filaments with CP bound to their distal free barbed ends. **(a)** Schematic depiction of the experimental strategy for studying twinfilin-mediated uncapping of CP-bound barbed ends of filaments anchored at their pointed ends. Actin filaments with free barbed end were polymerized from coverslip anchored spectrin-actin seeds by introducing 1 µM G-actin (15% Alexa-488 labeled) and 4 µM profilin in TIRF buffer. These filaments were then exposed to a flow containing 20 nM CP and subsequently allowed to age to ADP-F-actin for 15 minutes in presence of CP. The CP-capped filaments were then exposed to 1 µM G-actin (15% Alexa-488 labeled and 4 µM profilin alone (control) or additionally supplemented with 5 µM mTwf1 and filament uncapping was monitored. **(b)** Survival fraction of CP-bound ADP actin filaments as a function of time in absence or presence of 5 µM mTwf1. Experimental data (symbols) are fitted to a single-exponential decay function (lines) to determine CP dissociation rate k_-C_. Number of filaments analyzed for each condition: 60. **(c)** CP dissociation rate k_-C_ for ADP filaments in absence or presence of 5 µM mTwf1, determined from data in (b).

